# Identification, Genotyping, and Pathogenicity of *Trichosporon* spp. Isolated from Giant Pandas

**DOI:** 10.1101/386581

**Authors:** Xiaoping Ma, Yaozhang Jiang, Chengdong Wang, Yu Gu, Sanjie Cao, Xiaobo Huang, Yiping Wen, Qin Zhao, Rui Wu, Xintian Wen, Qigui Yan, Xinfeng Han, Zhicai Zuo, Junliang Deng, Zhihua Ren, Shumin Yu, Liuhong Shen, Zhijun Zhong, Guangneng Peng, Haifeng Liu, Ziyao Zhou

## Abstract

*Trichosporon* is the dominant genus of epidermal fungi in giant pandas and causes local and deep infections. To provide the information needed for the diagnosis and treatment of trichosporosis in giant pandas, the sequence of ITS, D1/D2, and IGS1 loci in 29 isolates of *Trichosporon* spp. which isolated from the body surface of giant pandas were combination to investigate interspecies identification and genotype. Morphological development was examined via slide culture. Additionally, mice were infected by skin inunction, intraperitoneal injection, and subcutaneous injection for evaluation of pathogenicity. The twenty-nine isolates of *Trichosporon* spp. were identified as belonging to 11 species, and *Trichosporon jirovecii* and *T. asteroides* were the commonest species. Four strains of *T. laibachii* and one strain of *T. moniliiforme* were found to be of novel genotypes, and *T. jirovecii* was identified to be genotype 1. *T. asteroides* had the same genotype which involved in disseminated trichosporosis. The morphological development processes of the *Trichosporon* spp. were clearly different, especially in the processes of single-spore development. Pathogenicity studies showed that 7 species damaged the liver and skin in mice, and their pathogenicity was stronger than other 4 species. *T. asteroides* had the strongest pathogenicity and might provoke invasive infection. The pathological characteristics of liver and skin infections caused by different *Trichosporon* spp. were similar. So it is necessary to identify the species of *Trichosporon* on the surface of giant panda. Combination of ITS, D1/D2, and IGS1 loci analysis, and morphological development process can effectively identify the genotype of *Trichosporon* spp.

## 1. Introduction

*Trichosporon* is a genus of fungi that belongs to the order *Tremellales* in the class *Tremellomycetes* (division *Basidiomycota*) and is widely distributed in nature(1). *Trichosporon* spp. can cause superficial fungal infections such as tinea pedis, onychomycosis, and dermoid infections(2). With the increasing prevalence of immunocompromised patients, the incidence of invasive fungal diseases has increased, and *Trichosporon* has become the second commonest genus of yeast fungus in deep fungal infections in patients with hematologic malignancies, granulocytic deficiency, and bone marrow transplants(3). Owing to difficulties in the classification and identification of *Trichosporon* spp., studies of invasive fungal infections have substantially lagged behind those of other species in areas such as clinical characteristics, antifungal susceptibilities, and the selection of therapeutic drugs, and this has become a difficult problem in the study of fungal infections(4).

*Trichosporon* spp. mainly cause skin and organ granuloma, and related pathogenicity studies have focused on *Trichosporon* spp. that colonize humans, such as *Trichosporon asahii, T. asteroides, T. inkin*, and *T. dermatis*. In recent years, reports of trichosporosis in animals have increased. Some cases in animals have been reported in the last decade, such as disseminated trichosporosis in cats(5), canine meningitis(6), and tortoise shell infection(7). A study described a case of systemic infection by *T. loubieri* in a cat with acute dyspnea, anorexia, and aggressive behavior; a cutaneous biopsy from ulcerated wounds revealed necrogranulomatous dermatitis and panniculitis with numerous intralesional fungal hyphae(5).

To enable accurate identification of *Trichosporon* spp., a number of molecular methods have been developed, of which DNA sequencing of the internal transcribed spacer (ITS) region, the D1/D2 domain of the 26S subunit of the rRNA gene region, and the intergenic spacer 1 (IGS1) region are the most frequently used. The IGS1 gene region has been particularly useful in phylogenetic studies and the description of intraspecies variation(8, 9). Ribeiro et al. identified 21 clinical isolates as belonging to six species on the basis of the ITS and IGS1 regions(10).

In 2009, Chagasneto et al. identified 22 isolates from human blood by analyzing the IGS1 region(11). In 2011, Guo identified 29 clinical isolates of *Trichosporon* by analyzing 3 loci, and eight *Trichosporon* spp. were found, of which *T. asahii* was the commonest(9). Identification of multiple *Trichosporon* spp. isolated from animals has not been reported. In addition, *Trichosporon* spp. are the dominant fungal species on panda skin(12). The identification at the species level is important for preventing dermatomycoses in giant pandas and improving related research.

At present, there have been few studies of the morphology of *Trichosporon* spp. In 2005, Li performed slide culture of six clinically common *Trichosporon* spp. and found no significant differences in colony morphology(13).

We collected 29 isolates of *Trichosporon* spp. from the skin of giant pandas at the China Conservation and Research Center for Giant Pandas, Ya’an. Owing to the shortage of sequence data for the IGS1 region in individual species, we had to analyze the ITS region, D1/D2 domain, and IGS1 region of the isolates to obtain accurate classification information. Afterward, the morphological development process was observed by the slide culture method. Mice were artificially infected, and their livers and skin were taken for pathological analysis.

## 2. Materials and methods

### 2.1 Sampling procedure

Samples were collected from clinically healthy giant pandas (22 females and 22 males) at the China Conservation and Research Center for Giant Pandas (Ya’an, China) in 2015–2016. Pandas that had been treated with antifungal drugs during the previous 6 months or with a recent history of disease were excluded from this study. The pandas lived in a semi-captive semi-enclosed breeding environment, were fed a diet of about 10% steamed cornbread and fruits and 90% bamboo shoots, and were allowed to drink water *ad libitum*(14).

All personnel involved in sampling wore sterile protective clothing, hats, masks, and latex gloves. When a panda ate fruits, 70% alcohol was used to sterilize the surface of the dorsal forearm, and then a suitable amount of dander was collected using sterilized surgical blades, scissors, and forceps. All samples were quickly placed in sterilized plastic sample bags, transported to the laboratory on ice within 2 h, and then immediately processed in a BSL-2 safety cabinet. No repeat sampling was performed on the same panda, and all 44 samples were processed with isolated *Trichosporon* spp.(14).

### 2.2 Fungal culture

Samples were streak-inoculated under aerobic conditions onto Sabouraud dextrose agar (SDA) (MOLTOX, Inc., Boone, NC) containing 4% (m/v) glucose, 1% (m/v) peptone, and 1.5% (m/v) agar. All media were supplemented with the antibiotics cycloheximide (0.05%, m/v) and chloramphenicol (0.005%, m/v).

Fungal culture was carried out in a BSL-2 safety cabinet in a bioclean room. Sterilized sealing film was used to cover each plate. Each sample was plated onto three culture plates with three control plates. All culture dishes were inoculated and stored at 25 °C for 7–30 days before being considered negative(14).

### 2.3 Molecular identification

Fungal DNA was extracted from pure culture isolates as described previously(15). Amplification of the ITS region, D1/D2 domain, and IGS1 region was performed as described with the primer pairs ITS1/ITS4 (ITS1: 5’-TCCGTAGGTGAACCTGCGG-3’; ITS4: 5’-TCCTCCGCTTATTGATATGC-3’), F63/R635 (F63: 5’-GCATATCAATAAGCGGAGCAAAAG-3’; R635: 5’-GGTCCGTGTTTCAAGACG-3’), and 26SF/5SR (26SF: 5’-ATCCTTTGCAGACGACTTGA-3’; 5SR: 5’-AGCTTGACTTCGCAGATCGG-3’), respectively^(8, 16)^. PCR amplification was performed in a 50 μl reaction mixture containing 19 μl 2 × Taq Master Mix (Tsingke Biotech Co., Ltd., Chengdu, China), 2 μl primers, 25 μl double-distilled water, and 2 μl fungal genomic DNA. The thermocycling conditions were as follows: 5 min at 98 °C (initial denaturation), 35 cycles of 10 s at 98 °C, 10 s at 58 °C, and extension at 72 °C for 10 s, and final extension for 4 min at 72 °C. A total of 8 μl of the amplified PCR products were visualized on 2% agarose gel after staining with GreenView (Solarbio, Beijing). The PCR products were then sequenced by Tsingke Biotech Co., Ltd. (Chengdu, China).

All the chromatograms of DNA sequences were examined to ensure high-quality sequences. For species identification, the sequences of the ITS region, D1/D2 domain, and IGS1 region were queried against the NCBI database (https://www.ncbi.nlm.nih.gov). The sequence of each locus and concatenated sequences were then aligned using the NCBI BLAST and formed the consensus sequences for all 29 isolates. Phylogenetic trees were computed with MEGA version 6 (Molecular Evolutionary Genetic Analysis software version 6.0.2; http://www.megasoftware.net) using the neighbor-joining method, in which all positions containing gaps and missing data were eliminated from the dataset. The ITS region plus the D1/D2 domain and IGS1 region were used to produce two separate phylogenetic trees. All sequences of the three genes from the 29 isolates were deposited in the GenBank database (https://www.ncbi.nlm.nih.gov/genbank/) and were assigned ID numbers (Table 1).

**Table 1.**
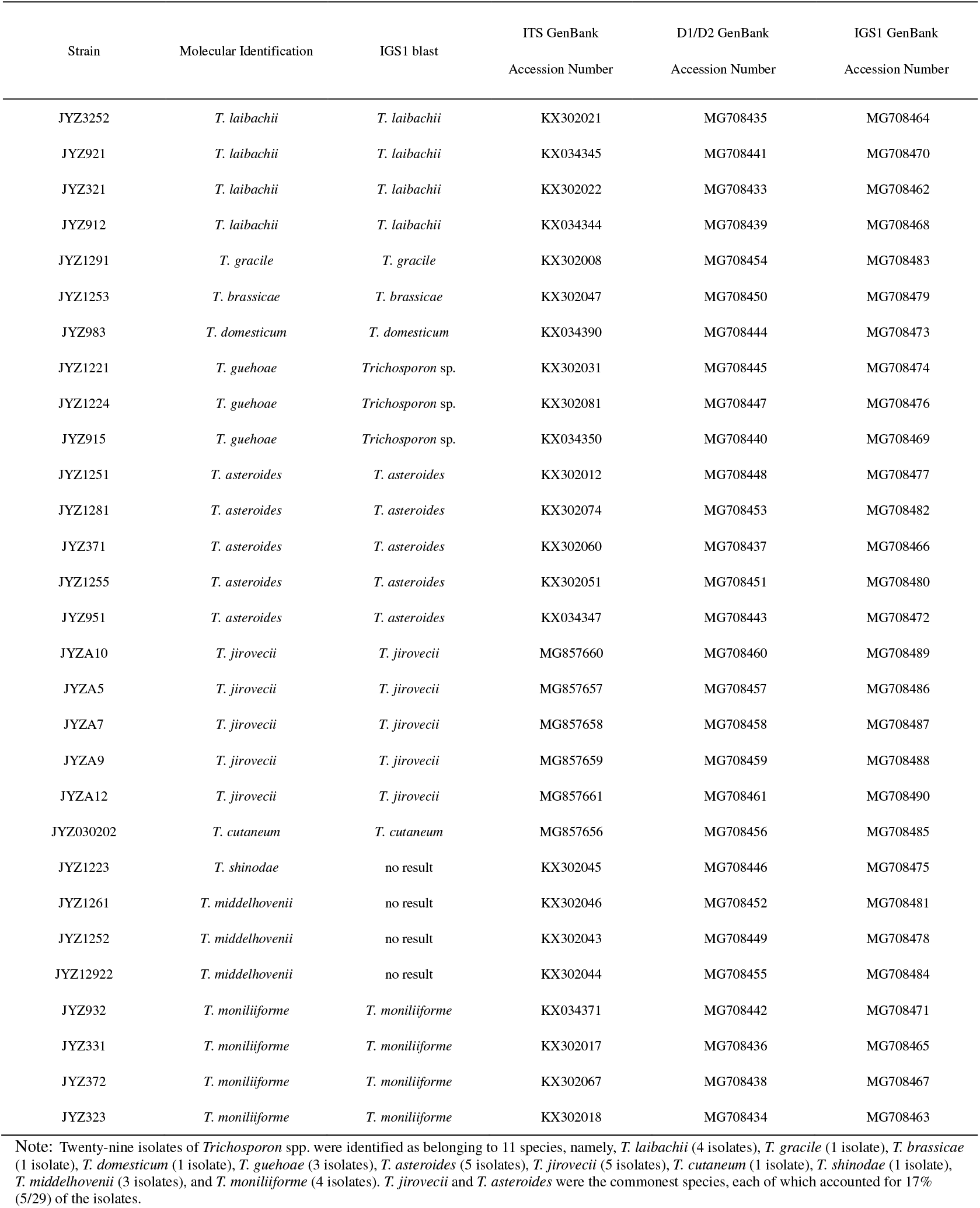
Nucleotide sequence accession numbers.

| Strain | Molecular Identification | IGS1 blast | ITS GenBank | D1/D2 GenBank | IGS1 GenBank |
| --- | --- | --- | --- | --- | --- |
|  |  |  | Accession Number | Accession Number | Accession Number |
| JYZ3252 | *T. laibachii* | *T. laibachii* | KX302021 | MG708435 | MG708464 |
| JYZ921 | *T. laibachii* | *T. laibachii* | KX034345 | MG708441 | MG708470 |
| JYZ321 | *T. laibachii* | *T. laibachii* | KX302022 | MG708433 | MG708462 |
| JYZ912 | *T. laibachii* | *T. laibachii* | KX034344 | MG708439 | MG708468 |
| JYZ1291 | *T. gracile* | *T. gracile* | KX302008 | MG708454 | MG708483 |
| JYZ1253 | *T. brassicae* | *T. brassicae* | KX302047 | MG708450 | MG708479 |
| JYZ983 | *T. domesticum* | *T. domesticum* | KX034390 | MG708444 | MG708473 |
| JYZ1221 | *T. guehoae* | *Trichosporon* sp. | KX302031 | MG708445 | MG708474 |
| JYZ1224 | *T. guehoae* | *Trichosporon* sp. | KX302081 | MG708447 | MG708476 |
| JYZ915 | *T. guehoae* | *Trichosporon* sp. | KX034350 | MG708440 | MG708469 |
| JYZ1251 | *T. asteroides* | *T. asteroides* | KX302012 | MG708448 | MG708477 |
| JYZ1281 | *T. asteroides* | *T. asteroides* | KX302074 | MG708453 | MG708482 |
| JYZ371 | *T. asteroides* | *T. asteroides* | KX302060 | MG708437 | MG708466 |
| JYZ1255 | *T. asteroides* | *T. asteroides* | KX302051 | MG708451 | MG708480 |
| JYZ951 | *T. asteroides* | *T. asteroides* | KX034347 | MG708443 | MG708472 |
| JYZA10 | *T. jirovecii* | *T. jirovecii* | MG857660 | MG708460 | MG708489 |
| JYZA5 | *T. jirovecii* | *T. jirovecii* | MG857657 | MG708457 | MG708486 |
| JYZA7 | *T. jirovecii* | *T. jirovecii* | MG857658 | MG708458 | MG708487 |
| JYZA9 | *T. jirovecii* | *T. jirovecii* | MG857659 | MG708459 | MG708488 |
| JYZA12 | *T. jirovecii* | *T. jirovecii* | MG857661 | MG708461 | MG708490 |
| JYZ030202 | *T. cutaneum* | *T. cutaneum* | MG857656 | MG708456 | MG708485 |
| JYZ1223 | *T. shinodae* | no result | KX302045 | MG708446 | MG708475 |
| JYZ1261 | *T. middelhovenii* | no result | KX302046 | MG708452 | MG708481 |
| JYZ1252 | *T. middelhovenii* | no result | KX302043 | MG708449 | MG708478 |
| JYZ12922 | *T. middelhovenii* | no result | KX302044 | MG708455 | MG708484 |
| JYZ932 | *T. moniliiforme* | *T. moniliiforme* | KX034371 | MG708442 | MG708471 |
| JYZ331 | *T. moniliiforme* | *T. moniliiforme* | KX302017 | MG708436 | MG708465 |
| JYZ372 | *T. moniliiforme* | *T. moniliiforme* | KX302067 | MG708438 | MG708467 |
| JYZ323 | *T. moniliiforme* | *T. moniliiforme* | KX302018 | MG708434 | MG708463 |

Note: Twenty-nine isolates of *Trichosporon* spp. were identified as belonging to 11 species, namely, *T. laibachii* (4 isolates), *T. gracile* (1 isolate), *T. brassicae* (1 isolate), *T. domesticum* (1 isolate), *T. guehoae* (3 isolates), *T. asteroides* (5 isolates), *T. jirovecii* (5 isolates), *T. cutaneum* (1 isolate), *T. shinodae* (1 isolate), *T. middelhovenii* (3 isolates), and *T. moniliiforme* (4 isolates). *T. jirovecii* and *T. asteroides* were the commonest species, each of which accounted for 17% (5/29) of the isolates.

### 2.4 Morphological development process

All 29 isolates were identified as *Trichosporon* spp. after molecular identification. Microscopic observations of the 29 isolates were made after slide culture on SDA(1). A 0.5 ml sample of melted medium was injected into a closed glass Petri dish, which comprised a slide glass, a cover glass, and a copper ring with a hole in the wall, and was inoculated via the hole(17, 18). All isolates were incubated at 25 °C and were observed after 24, 48, 72 and 96 h. The cover glass was stained with 5 ml cotton blue (Hopebio, Qingdao) and was observed with a microscope (BX51, Olympus).

### 2.5 Pathogenicity experiment

#### 2.5.1 Animal experiment

Each *Trichosporon* sp. was inoculated by skin inunction (We abraded skin with emery paper until a slight bleedingand cut the hair (2 cm×2 cm) on the back of mouse after anesthesia by diethyl ether), subcutaneous injection, and intraperitoneal injection into immunosuppressed and non-immunosuppressed mice. In total, six groups were used, each of which comprised three mice. Groups A (intraperitoneal injection), B (subcutaneous injection), and C (skin inunction) were immunosuppressed (Mice were given intraperitoneal injection with 50 mg/kg cyclophosphamide at intervals of 48 hours, three times in total, and each mouse was given 15 mg of penicillin sodium under the skin.); groups D (intraperitoneal injection), E (subcutaneous injection), and F (skin inunction) were non-immunosuppressed. In totalof 216 sex-matched SPF Kunming mice (Dashuo Science and Technology Co., Ltd., China), which belonged to 11 experiment groups and one control group, with ages of 6–8 weeks were used.

#### 2.5.2 Preparation of fungal suspension and inoculation

Before being inoculated, mice in the immunosuppressed groups were intraperitoneally injected with *Trichosporon* spp. that had been cultured in SDA at 25 °C for 5 d. The mycelium and spores were scraped, washed with physiological saline, and mixed well. A hemocytometer was used to adjust the concentration of the spore suspension to 1 × 10^7^ CFU/ml. Except in the control group, each mouse was inoculated with 0.1 ml fungal suspension. Mice in the control group received 0.1 ml physiological saline instead. The backs of the mice treated by skin inunction were shaved, sterilized with 75% (w/w) alcohol, and lightly abraded with a 25G needle, and then 0.1 ml fungal suspension was gently rubbed onto the skin with a sterile injector.

#### 2.5.3 Tissue sample processing

The diet and clinical symptoms of the mice were observed daily. The mice were anesthetized with 5 ml diethyl ether (Chengdu Kelong Chemical Reagent Factory), decapitated, and dissected to observe lesions on the seventh day after infection. The livers of mice in groups A and D and skin lesions from mice in the other groups were taken for fungal culture and pathological evaluation. The livers and skin were placed in 100 ml 4% formalin (Chengdu Kelong Chemical Reagent Factory; 500 ml w/w) for histopathological study via staining with hematoxylin/eosin and periodic acid/Schiff stain(6).

## 3. Results

### 3.1 Molecular identification and genotyping

The interspecies identification of 22 isolates of *Trichosporon* spp. was performed from the data in Fig. 1. Seven strains of *Trichosporon* (JYZ1221, JYZ1224, JYZ915, JYZ1223, JYZ1261, JYZ1252, and JYZ12922) could not be identified because of the lack of sequence information for IGS1 in the GenBank database. However, it was determined that these seven *Trichosporon* strains belonged to three species. The interspecies identification of all 29 isolates of *Trichosporon* spp. was performed using the data in Fig. 2. The structures of the two phylogenetic trees were basically identical, and the method could be used to determine the accuracy of the identification of the 29 isolates. The 29 isolates of *Trichosporon* spp. were identified as belonging to 11 species, namely, *T. laibachii* (4 strains), *T. gracile* (1), *T. brassicae* (1), *T. domesticum* (1), *T. guehoae* (3), *T. asteroides* (5), *T. jirovecii* (5), *T. cutaneum* (1), *T. shinodae* (1), *T. middelhovenii* (3), and *T. moniliiforme* (4). *T. jirovecii* and *T. asteroides* were the commonest species (17%, 5/29). Moreover, *T. middelhovenii* and *T. shinodae* were isolated from the surfaces of animals for the first time.

**Fig. 1.**
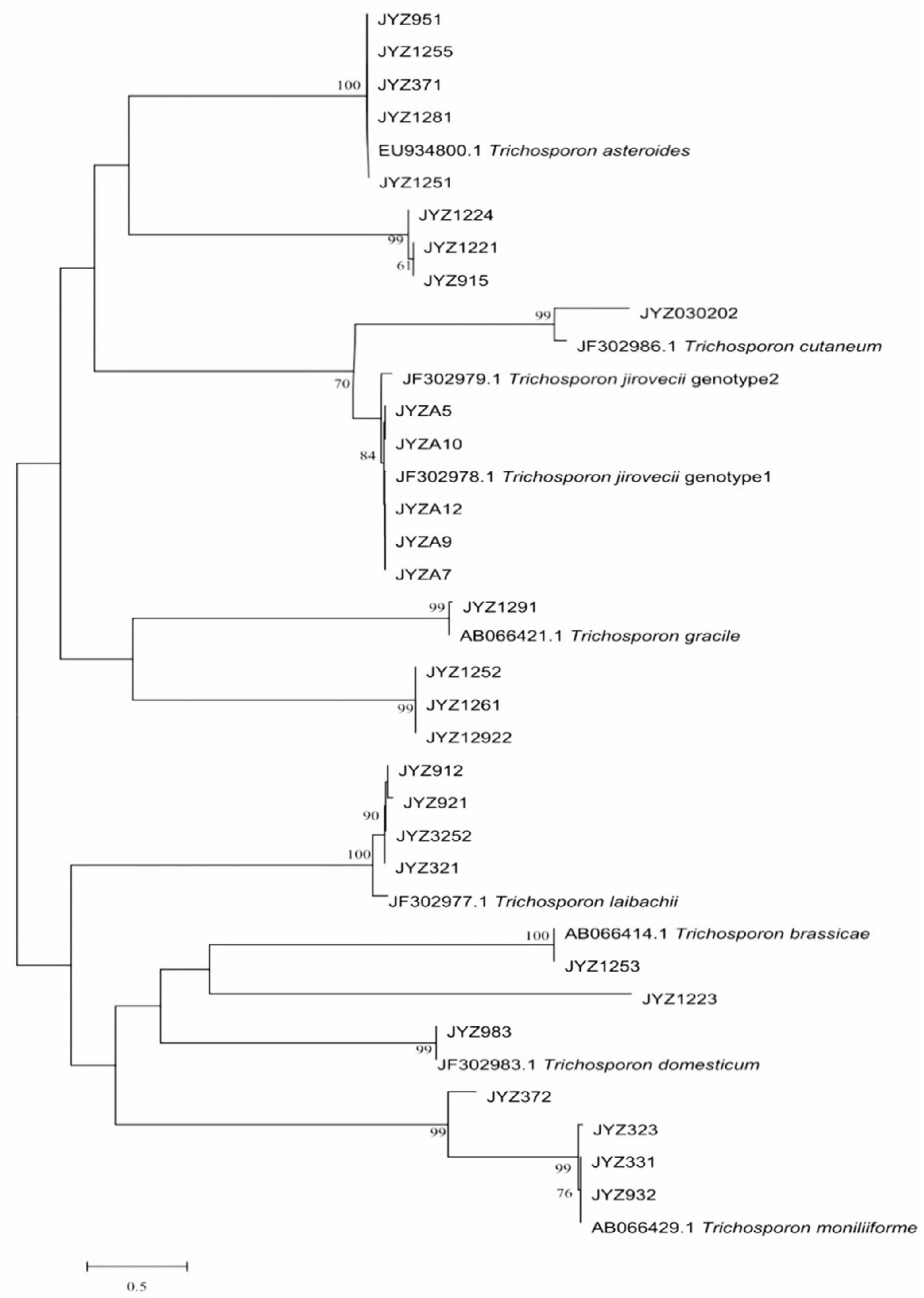
Phylogenetic tree based on IGS1 sequences

**Fig. 2.**
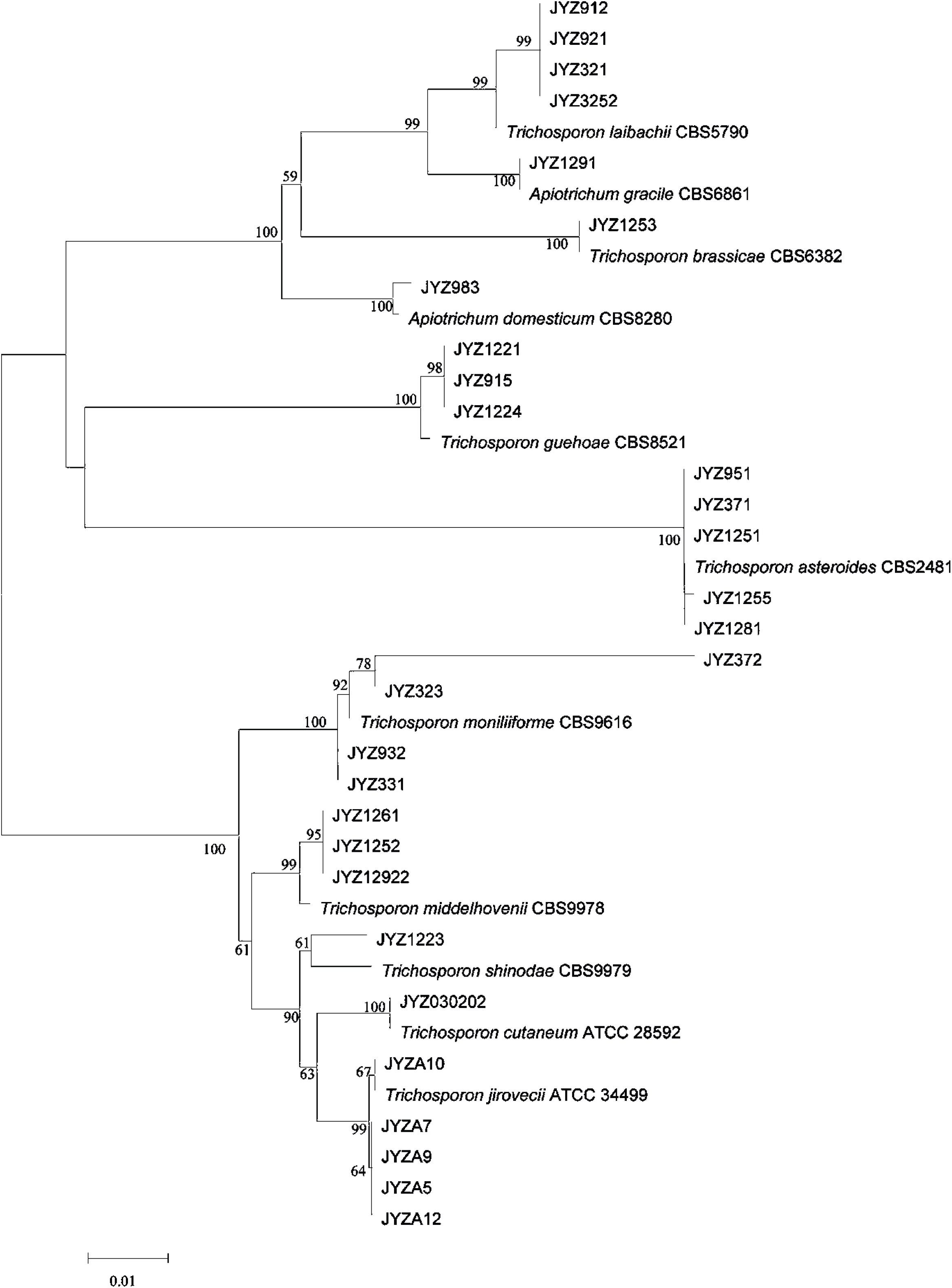
Phylogenetic tree based on ITS sequences plus D1/D2 sequences

In this study, preliminary genotyping was performed on *Trichosporon* spp. *T. asteroides* (JYZ1251, JYZ1281, JYZ371, JYZ1255, and JYZ951) had the same genotype as its reference strain, which was isolated from the blood of immunocompromised patients(11), whereas *T. laibachii* (JYZ3252, JYZ921, JYZ321, and JYZ912) had different genotypes. The reference strain was isolated from humans. The five isolates of *T. jirovecii* were identified as having genotype 1(9). *T. brassicae* (JYZ1253) had the same genotype as the reference strain, which was isolated from cabbage(8). *T. gracile* (JYZ1291) had the same genotype as the reference strain, which was isolated from spoiled milk(8). *T. domesticum* (JYZ983) and the reference strain had the same genotype; the reference strain was isolated from human sputum(8). Three strains of *T. moniliiforme* (JYZ932, JYZ331, and JYZ323) had the same genotype as the reference strain, which was isolated from curdling milk(8), but the strain JYZ372 did not have this genotype. It was difficult to determine whether *T. cutaneum* had the same genotype as the reference strain. According to the phylogenetic tree, the genetic relationship was distant and it may not have had the same genotype.

### 3.2 Morphological development process

The morphological development processes of the *Trichosporon* spp. were clearly different, and the difference was significant for the processes of single-spore development. Spores of *T. moniliiforme* and *T. shinodae* tended to be self-replicating at the beginning of their development and reproduced mainly by budding. Spores of the other species tended to become mycelia and reproduced mainly by the differentiation of mycelia to form spores or produce arthrospores. Morphological development could be used as an important basis for the identification of *Trichosporon* spp. For example, *T. moniliiforme* produced a large number of spores, and the spores were transformed into an oval shape after development was completed. The growth of *T. shinodae* was the slowest among the *Trichosporon* spp.; the shape of mycelia was specific and served as a basis for identification. *T. laibachii* produced a large number of arthrospores and had the unique feature that the mycelia were folded together. The main method of growth of *T. guehoae* comprised the reproduction of arthrospores by budding, and it had the specific feature that new mycelia grew from gaps in segmented mycelia. The morphological appearances of *T. brassicae*, *T. domesticum* and *T. gracile* were analogous in some ways. All these *Trichosporon* spp. tended to reproduce via the differentiation of mycelia into spores, and few arthrospores were seen during the process of development. The shapes of spores that differentiated from mycelia were different: those of *T. gracile* were quadrilateral, those of *T. domesticum* were round, and those of *T. brassicae* were disciform. The arthrospores of *T. middelhovenii* were fusiform and were always located in a bifurcation of the mycelium. This characteristic was distinct from other *Trichosporon* spp. *T. asteroides* had the feature that the spores overlapped each other, and the mycelia were thin and short. The mycelia could differentiate into spores, and the pigmentation of the spores was uneven, as shown by dyeing with cotton blue. The morphologies of *T. jirovecii* and *T. cutaneum* were similar, but the mycelia of *T. cutaneum* were more curved and tended to differentiate into spores, in contrast to *T. jirovecii. Trichosporon* spp. have individual morphological characteristics and hence could be distinguished by means of a comparison of their morphological development processes.

After culture for 24 h, spores of *T. moniliiforme* divided independently (Fig. 3A), no hyphae differentiated into spores (Fig. 3B, C, and D), and the mycelium and spores were evenly stained (Fig. 3B, C, and D). The mycelium produced a large number of arthrospores (Fig. 3C), and the shape of the spores changed from circular (Fig. 3C) to elliptical (Fig. 3D).

**Fig. 3.**
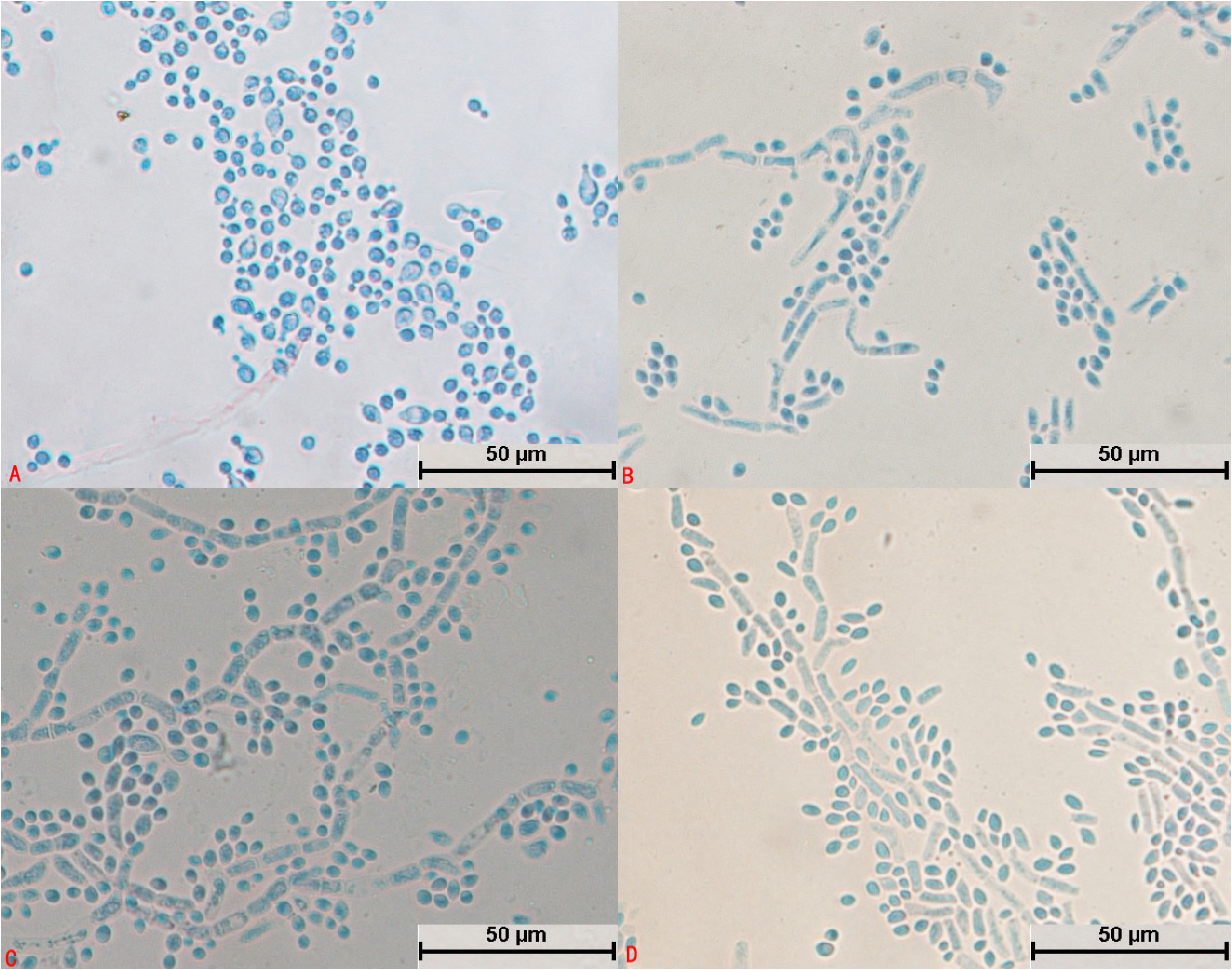
Morphological development process of *Trichosporon moniliiforme* from day 1 to day 4. A: Most spores divided and a small number of spores expanded; B: Scattered hyphae appeared and began to produce conidia on day 2; C: Basic hyphae formed; D: Spores increased in number and their shapes were transformed from circular to elliptical.

The spores of *T. laibachii* grew into mycelium after cultivation for 24 h, and no spores divided independently (Fig. 4A). The mycelium spread radially (Fig. 4A) and was folded together (Fig. 4C), and differentiated into spores. Arthrospores were abundant, whereas spores were round and few in number (Fig. 4B, C, and D). The spores and mycelium were unevenly colored: the spores were darker, whereas the mycelium was lighter (Fig. 4B, C, and D). Hyphal folding was typical in structure.

**Fig. 4.**
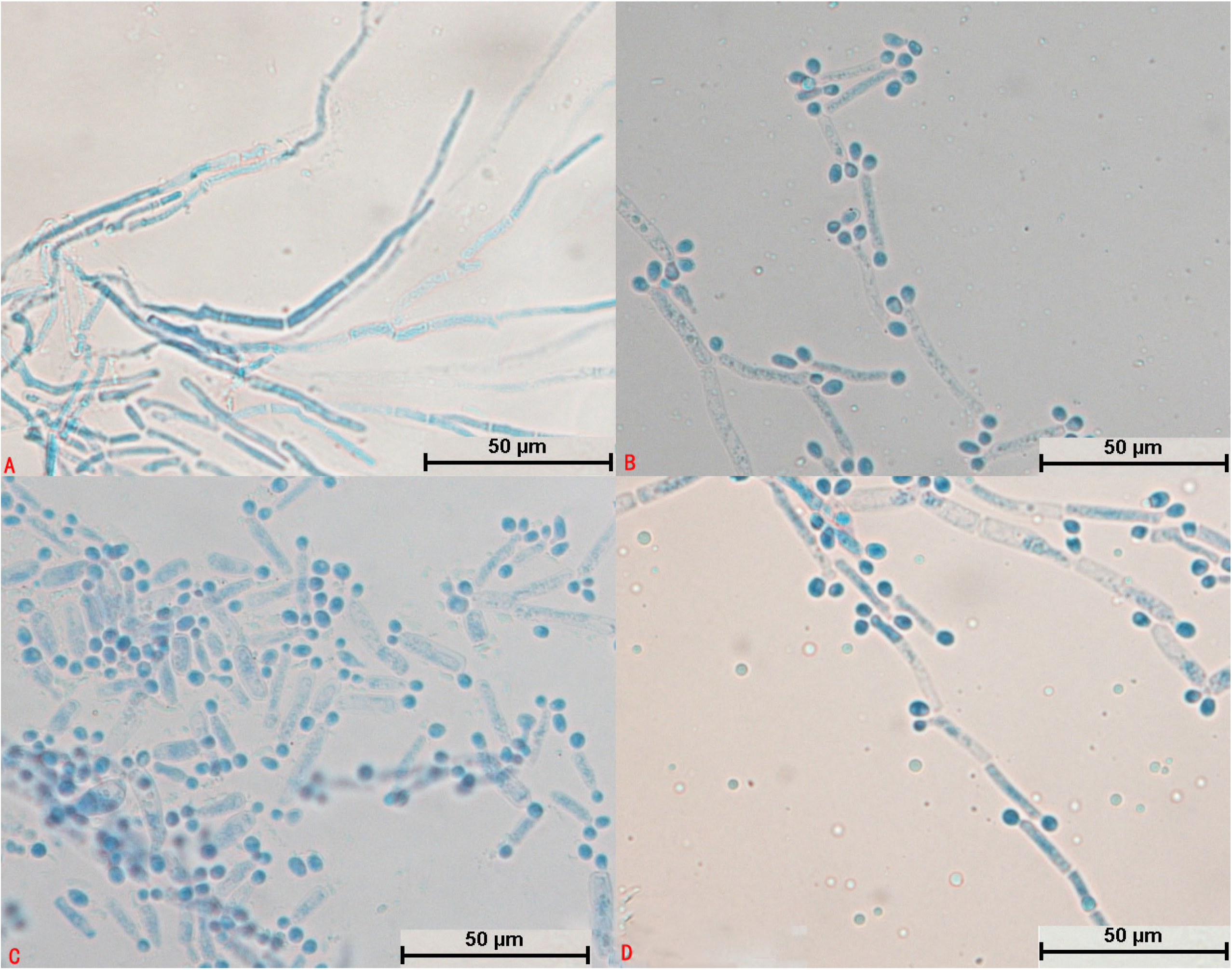
Morphological development process of *Trichosporon laibachii* from day 1 to day 4. A: Some spores germinated and mycelium began to divide; B: Mycelium produced arthrospores; C: Mycelium was folded; D: Mature mycelium and arthrospores.

The structural development of *T. guehoae* was completed after cultivation for 24 h (Fig. 5A). The mycelium spread radially but was scattered (Fig. 5A, B, C, and D). A large number of round spores were produced as grape-like clusters (Fig. 5A, B, and C). Hyphae and spores were evenly stained (Fig. 5A, B, C, and D). The mycelium grew from a segmented section to form a new mycelium (Fig. 5B), and the new spores were generated by arthrospores (Fig. 5D); this was a typical structure.

**Fig. 5.**
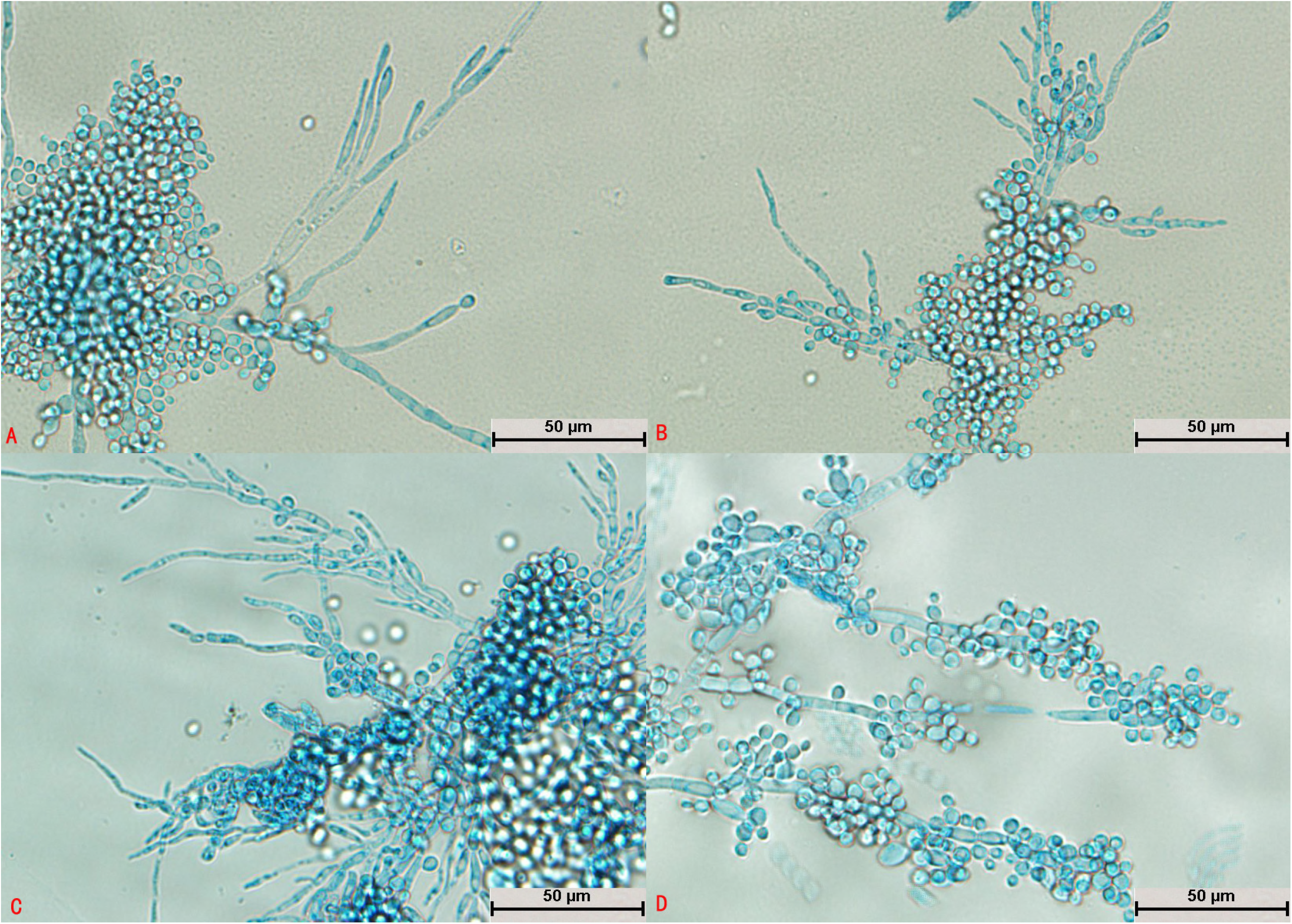
Morphological development process of *Trichosporon guehoae* from day 1 to day 4. A: Mycelial buds formed new mycelium; B: New mycelium appeared at the point at which hyphae were segmented; C: A large number of botryoidal spores of *T. guehoae* were observed; D: New spores developed from arthrospores.

After *T. gracile* was cultured for 24 h, a large amount of hyphae appeared and became segmented, and no spores could be seen (Fig. 6A). It could be seen that the mycelium differentiated into spores (Fig. 6B, C and D). A large number of spores were produced, which were mostly square (Fig. 6C) and became oval after reaching maturity (Fig. 6D). The mycelia and spores were evenly stained (Fig. 6C). Some hyphae did not divide into sections and differentiated into spores at intervals; this was a typical structure (Fig. 6D).

After culture for 24 h, the spores of *T. domesticum* swelled to form mycelia and no spores divided independently (Fig. 7A). Hyphae were abundant and parallel to each other and had spindle-type buds (Fig. 7B). A large number of hyphae differentiated into spores (Fig. 7C and D). The number of spores was small with no arthrospores, and the spores were round (Fig. 7B, C, and D). The mycelia and spores were evenly stained (Fig. 7A, B, C, and D).

**Fig. 6.**
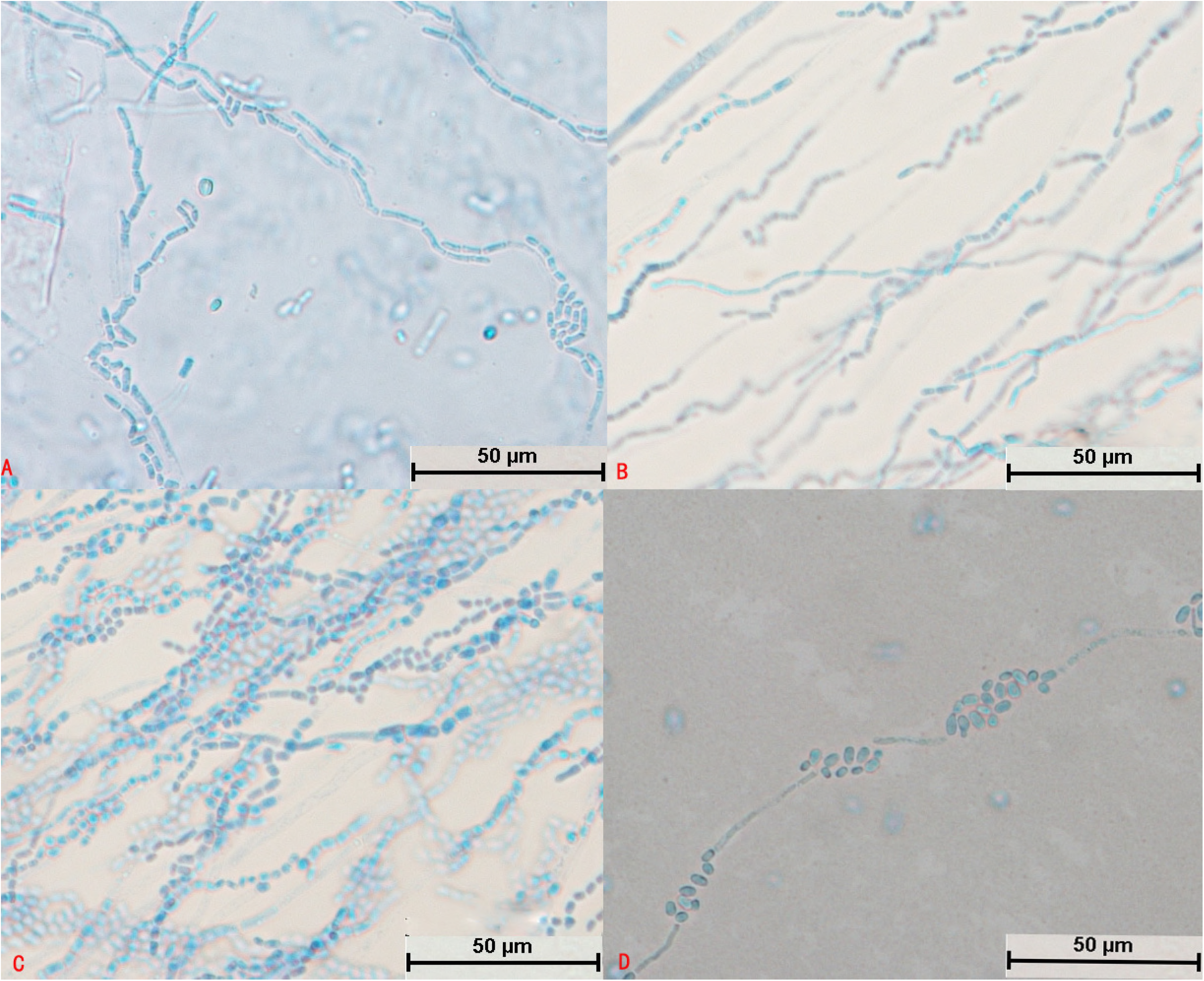
Morphological development process of *Trichosporon gracile* from day 1 to day 4. A and B: Hyphae were segmented; C: A large number of hyphae differentiated into spores, some of which were square; D: Mycelium divided into oval spores.

**Fig. 7.**
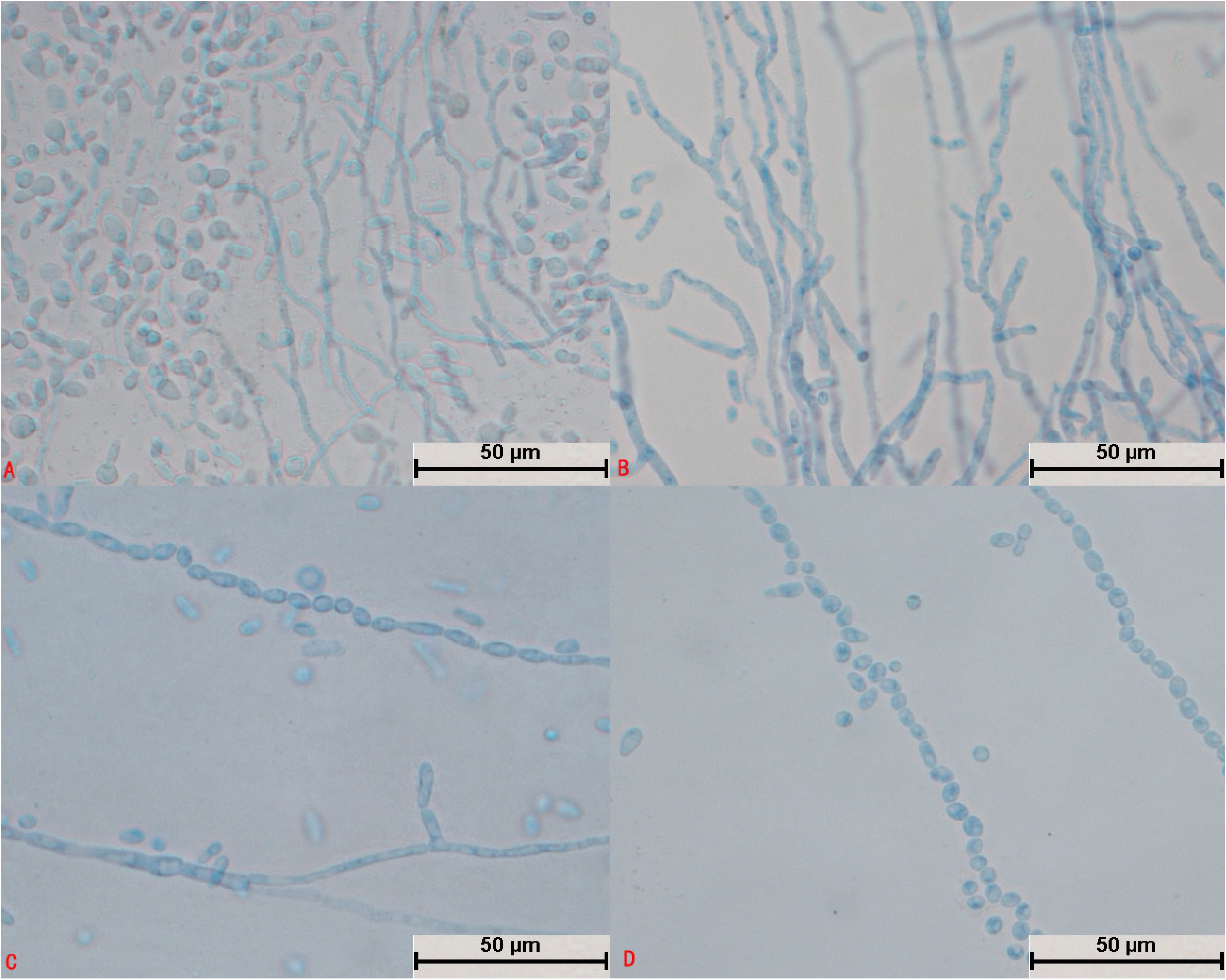
Morphological development process of *Trichosporon domesticum* from day 1 to day 4. A: Spores expanded and sprouted; B: Mycelium had multiple branches and spindle-shaped buds; C: Mycelium began to differentiate into oval spores; D: Mycelium had completely differentiated into oval spores.

After culture for 24 h, the spores of *T. brassicae* exhibited no significant changes (Fig. 8A), whereas after culture for 48 h the spores swelled and formed hyphae (Fig. 8B). No spores were found to divide independently (Fig. 8A and B). The mycelia became segmented and were distributed parallel to each other (Fig. 8C). It could be seen that the mycelium differentiated into spindle-shaped spores (Fig. 8D). The mycelium and spores were evenly stained (Fig. 8D), and there were few arthrospores (Fig. 8C).

**Fig. 8.**
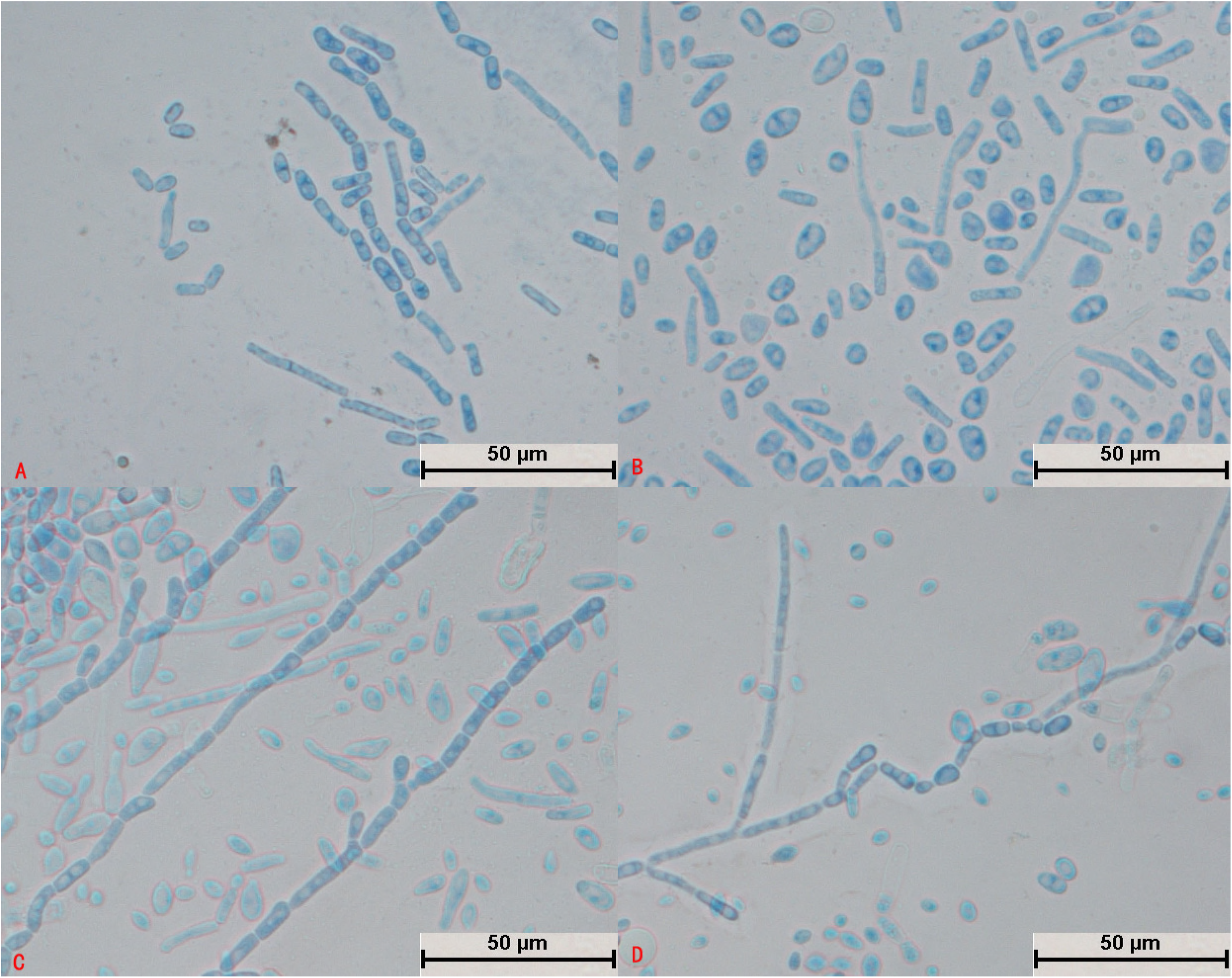
Morphological development process of *Trichosporon brassicae* from day 1 to day 4. A: Spindle-shaped spores; B: Spores sprouted; C: Hyphae were segmented and spores were less oval; in the segmented hyphae, the arthrospores were fewer in number and spindle-shaped, and some spores developed into mycelium; D: Hyphae differentiated into spindle-shaped spores.

After cultivation for 24 h, some spores of *T. shinodae* swelled (Fig. 9A), whereas other spores divided independently (Fig. 9B). The hyphae were short and sparse and grew very slowly (Fig. 9C). A large number of round arthrospores were produced (Fig. 9D). The spores and mycelium were evenly colored (Fig. 9C and D). Crude short mycelium was produced after culture for 72 h (Fig. 9C); this was a typical structure.

**Fig. 9.**
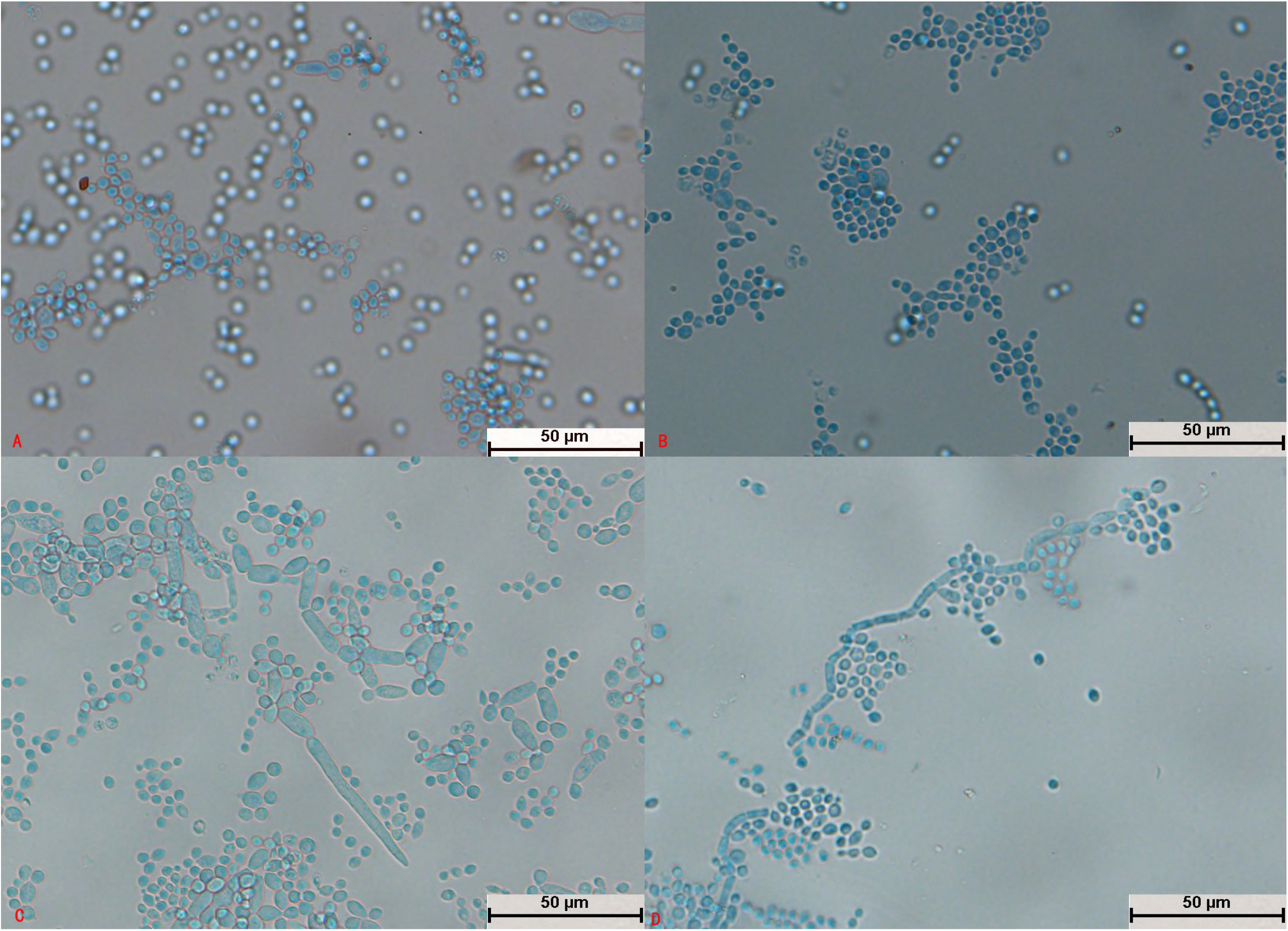
Morphological development process of *Trichosporon shinodae* from day 1 to day 4. A: Spores swelled and budded; B: Spores divided; C: Spores formed thick and short hyphae; D: Mycelium produced a large number of arthrospores.

After culture for 24 h, *T. asteroides* formed slender hyphae, and no spores divided independently (Fig. 10A). The mycelium was elongated (Fig. 10C) and could differentiate into spores (Fig. 10B and D). Spores on bifurcated mycelium aggregated into spheres (Fig. 10C); a large number of spores were produced, and the spores were round, oval, or spindle-shaped (Fig. 10B and D). The spores were not evenly pigmented and some were dark in color (Fig. 10D). The bifurcation of the mycelia and the aggregation of spores into spheres were typical structures (Fig. 10C).

**Fig. 10.**
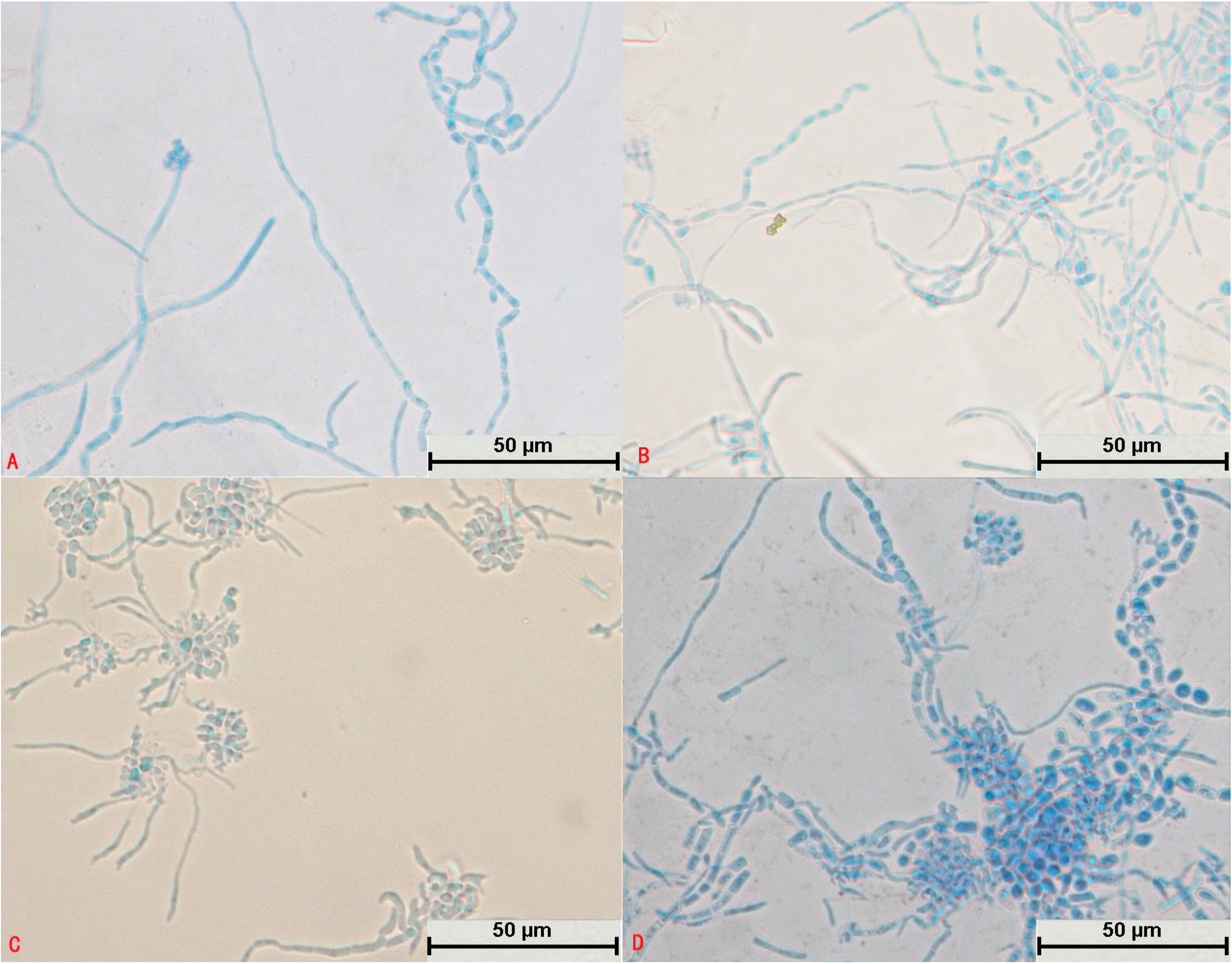
Morphological development process of *Trichosporon asteroides* from day 1 to day 4. A: Tenuous mycelium, some of which began to become segmented; B: Mycelium differentiated into circular, oval, and spindle-shaped spores; C: The tail ends of mycelium produced a large number of small, aggregating spores and bifurcated; D: Mycelium differentiated into darker spores.

After *T. middelhovenii* was cultivated for 24 h, hyphae were generated and no spores were found to divide independently (Fig. 11A). The hyphae were elongated (Fig. 11A and D), and their segments were small and indistinct (Fig. 11B and D). No hyphae differentiated into spores. Spores at bifurcations were spindle-shaped (Fig. 11A, B, C, and D). The spores were few in number and darker (Fig. 11C and D). Shuttle-type articular spores were characteristic structures (Fig. 11A, B, C, and D).

**Fig. 11.**
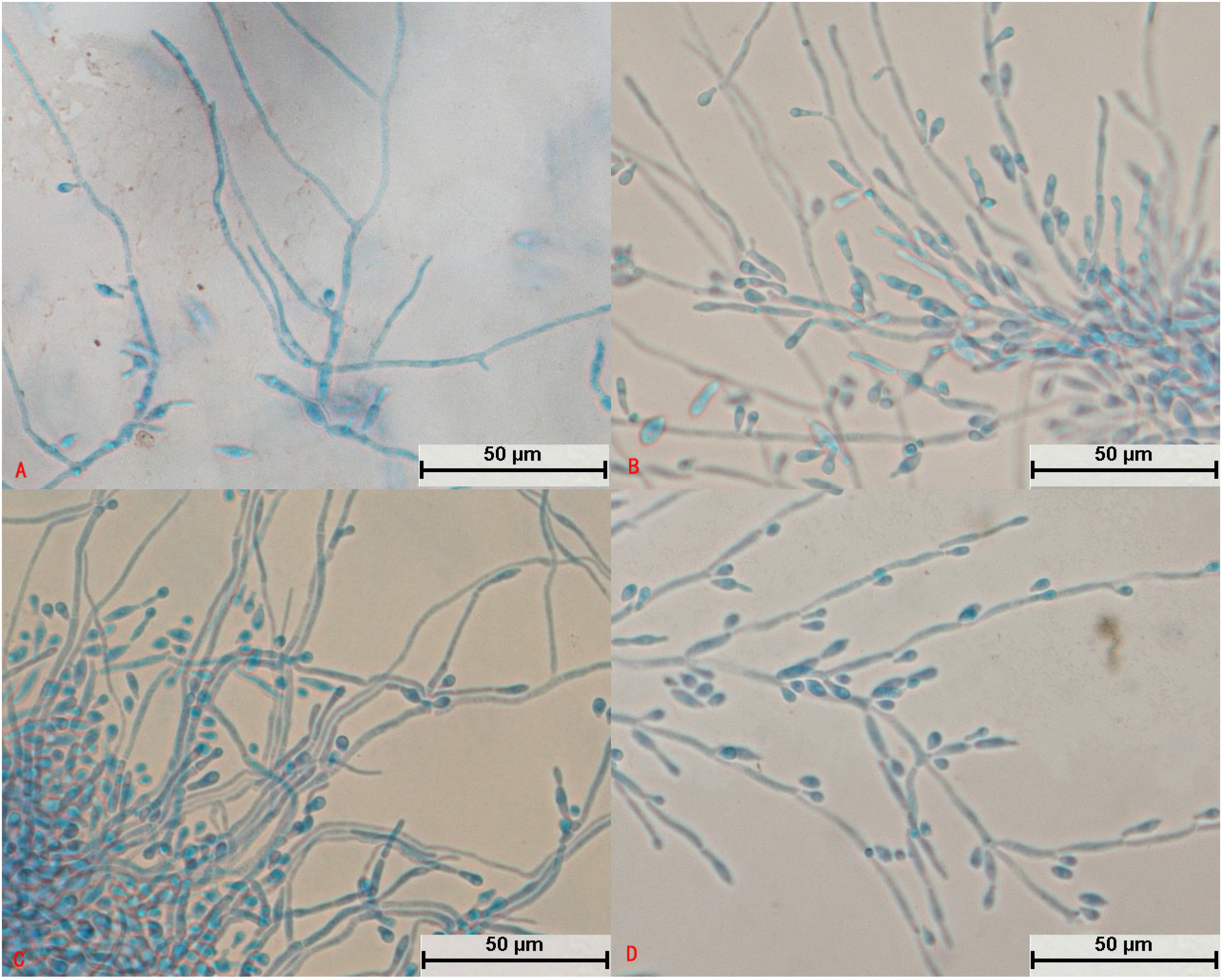
Morphological development process of *Trichosporon middelhovenii* from day 1 to day 4. A: Hyphae were slender and did not divide; B: Hyphae produced fusiform arthrospores; C: Arthrospores budded to form new hyphae; D: Mature mycelium produced a large number of fusiform spores.

After *T. jirovecii* was cultivated for 24 h, hyphae appeared and no spores were seen to divide independently (Fig. 12A). The hyphae were segmented and radial (Fig. 12A) and differentiated into spores (Fig. 12D). Arthrospores were round and large in number (Fig. 12B); mycelia and spores were uniformly colored (Fig. 12D).

**Fig. 12.**
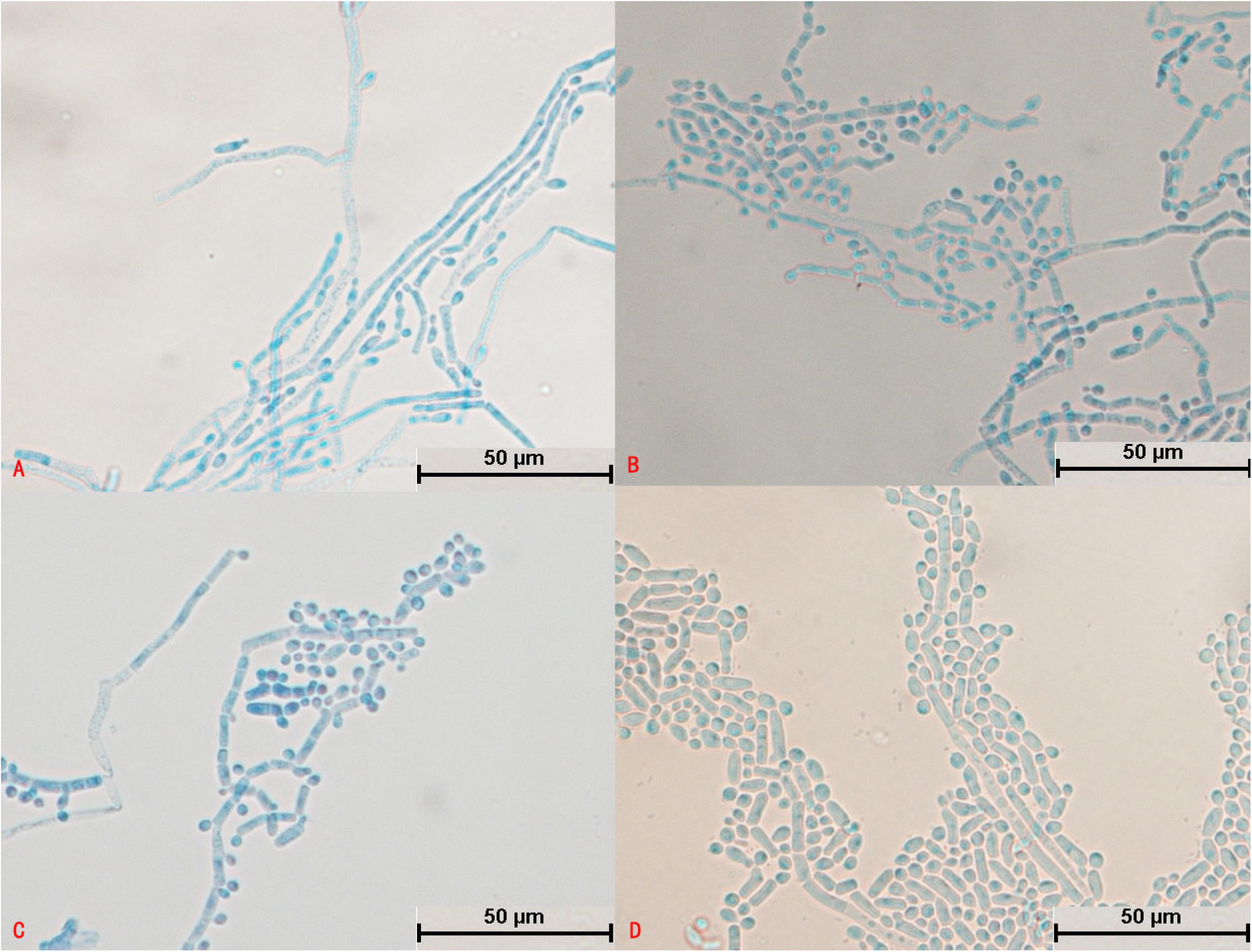
Morphological development process of *Trichosporon jirovecii* from day 1 to day 4. A: Mycelium was slightly segmented and formed arthrospores; B and C: Mycelium formed a large number of circular arthrospores; D: Mycelium differentiated into spores.

After cultivation for 24 h, spores of *T. cutaneum* swelled and budded, and some spores divided independently (Fig. 13A). The mycelium was curved and segmented (Fig. 13B and C) and differentiated into spores (Fig. 13D). A large number of round arthrospores (Fig. 13C) were produced. The mycelium and spores were evenly stained (Fig. 13D), and curved mycelium was a typical structure (Fig. 13B and D).

**Fig. 13.**
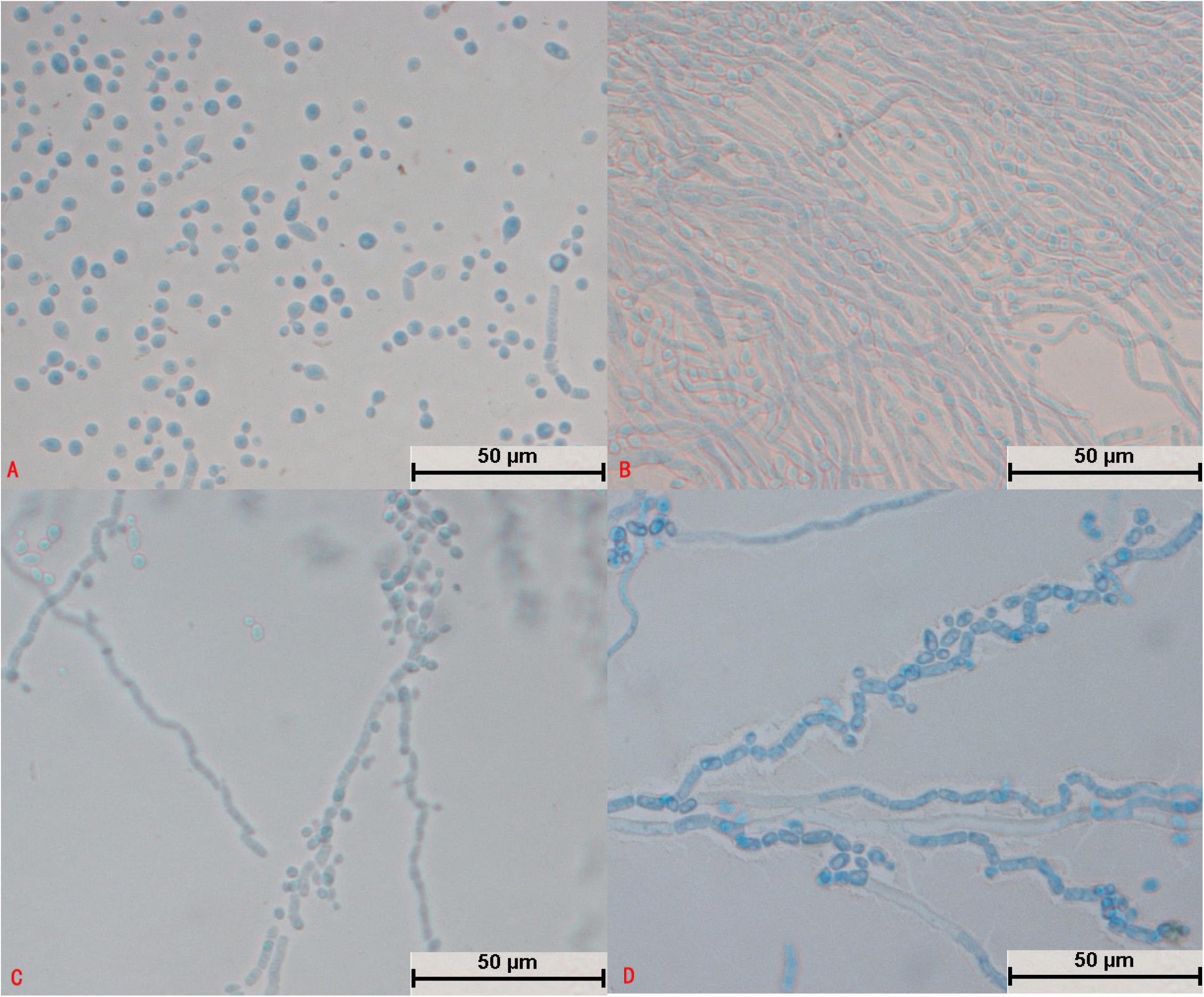
Morphological development process of *Trichosporon cutaneum* from day 1 to day 4. A: Some spores expanded and sprouted, and some spores divided; B: Spores formed curved mycelium; C: Most mycelium was segmented, and the mycelium produced round arthrospores; D: Mycelium began to fold and differentiated into spores.

### 3.3 Pathogenicity

The results of this study indicated that *Trichosporon* spp. mostly caused necrosis or swelling of hepatocytes and enlargement of the inter-hepatocyte space, and necrosis of hepatocytes mostly occurred near liver vessels. Subcutaneous injection of *Trichosporon* spp. caused lymphocyte infiltration into the skin, abscesses, and thickening of the stratum corneum. Mice that were inoculated via skin inunction had no obvious lesions, and most of them exhibited changes in the thickness of the stratum corneum, which in some cases resulted in subcutaneous abscesses. Tissues infected with *Trichosporon* spp. have a strong tendency to bleed, which causes congestion in both the skin and the liver. Most of the *Trichosporon* spp. caused significant damage to the liver and skin, for example, *T. laibachii, T. brassicae, T. guehoae, T. asteroides, T. jirovecii, T. cutaneum, T. shinodae*, and *T. middelhovenii. T. asteroides, T. laibachii, T. brassicae, T. guehoae, T. cutaneum, T. shinodae*, and *T. middelhovenii* all produced spores in the skin infection model (Fig.S1~S10). In particular, *T. asteroides* gave rise to disseminated infections in the reticular layer of the skin (Fig. 14G1) and budding in the dermis (Fig. 14G2). *T. gracile, T. moniliiforme*, and *T. domesticum* caused inconspicuous pathological changes, and hence their pathogenicity was weak.

**Fig. 14.**
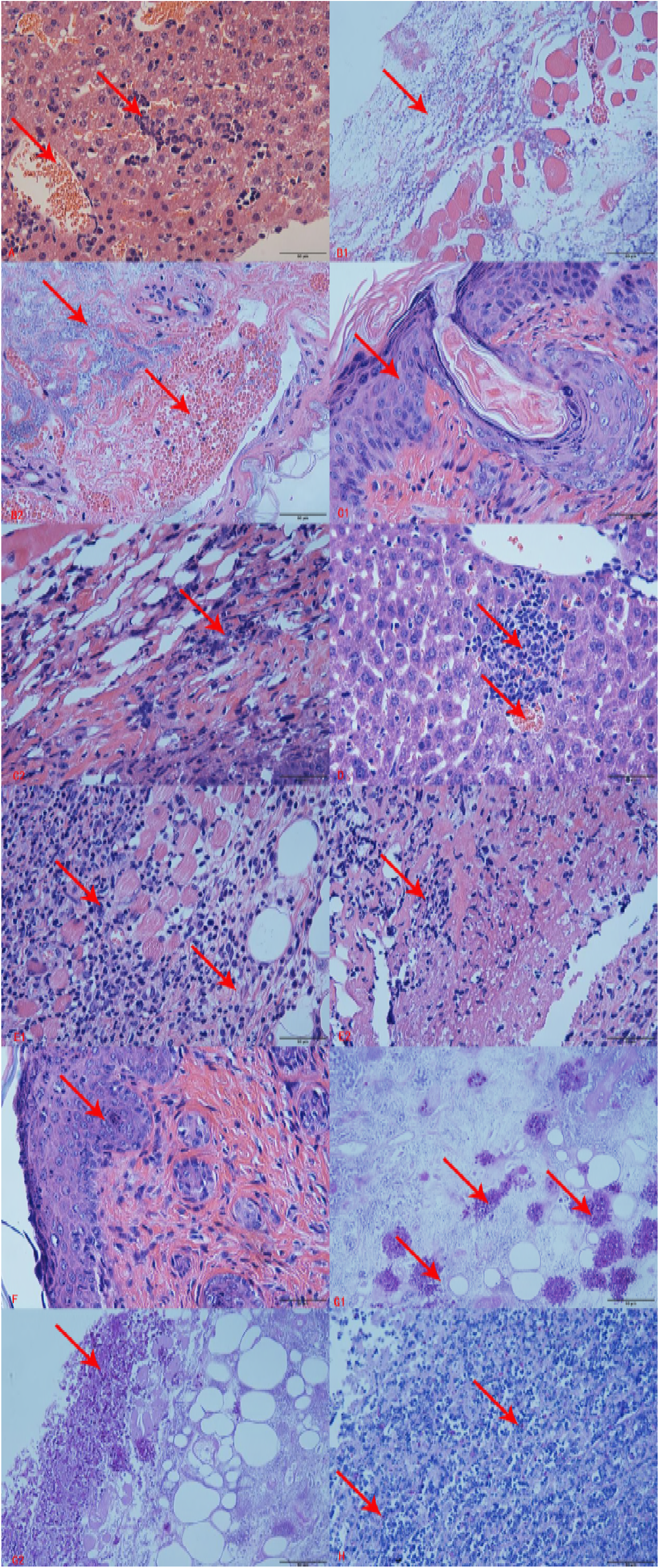
Pathological sections of tissue damaged by *Trichosporon asteroides* (JYZ1255) infection. A: Hepatocyte necrosis, hepatic sinusoidal congestion, and unclear hepatic cord structure; B1: Abscess of the dermis, necrosis of muscle tissue, and blood capillary congestion; B2: Epidermis with a large area of blood stasis and abscess; C1: Thickening of granular layer of skin and cuticle; C2: Infiltration of neutrophilic granulocytes into the epidermis; D: Hepatocyte necrosis and hepatic sinusoidal congestion; E1: A large number of neutrophils infiltrated into the reticular layer, muscle tissue, and dermis, and some cells are necrotic; E2: Osteonecrosis of the dermis and hyperplasia of connective tissue; F: Thickening of granular layer of skin and cuticle; G1: Spores stained with periodic acid/Schiff stain (PAS) in the reticular layer of the skin of a mouse in group B; G2: Spores stained with PAS in the dermis and spore germination in a mouse in group B; H: Spores stained with PAS in the dermis of a mouse in group E.

## 4. Discussion

### 4.1 Interspecies identification of *Trichosporon* spp.

There have been reports on the isolation and identification of fungi from the body surface of the giant panda(12). It was concluded that *Trichosporon* spp. were the dominant genus among skin flora on the giant panda(12). Recently, there have been many reports on infections by *T*. asahii(19-24), but few mentions of other *Trichosporon* spp.(6, 25). However, there have been reports that some animals are susceptible to rare *Trichosporon* spp.(25-27). Because the phylogenetic relationship between *Trichosporon* spp. was very close, it was impossible to distinguish the different species of *Trichosporon* spp. according to the ITS region or D1/D2 domain in every case(28). The sequence similarity between the ITS regions of *T. asahii* and *T. asteroides*, in which only two or three bases are different, is 99–99.3%, and *T. montevideense* and *T. domesticum* have identical ITS regions(29). Scorzetti *et al.* found that the differences between the 28s rDNA D1/D2 domains of different *Trichosporon* spp. are greater than those between the corresponding ITS regions. The ITS regions of *T. laibachii* and *T. multisporum* are identical, and seven bases are different in the D1/D2 domains. Two bases are different in the D1/D2 domains of *T. montevideense* and *T. domesticum*(30). Guo amplified all three loci (ITS, D1/D2, and IGS1) and constructed a phylogenetic tree for the ITS region and D1/D2 domain and a separate phylogenetic tree for the IGS1 region. Both trees could completely distinguish the *Trichosporon* spp.(9). In this study, seven strains could not be identified by their IGS1 regions because of the lack of sequence information for IGS1 in the NCBI database. Hence, we used the joint contribution of the ITS region and D1/D2 domain, which we compared with the phylogenetic tree for IGS1. It was found that the clades of the phylogenetic trees were basically identical and authenticated each other, so that all 29 strains could be identified completely.

### 4.2 Pathogenicity of dominant *Trichosporon* spp. isolated from pandas

*T. asteroides* and *T. jirovecii* were the dominant *Trichosporon* spp. that were isolated from the giant panda samples, and these species are widely present in giant pandas(12). Their pathogenicity has a great influence on the health of giant pandas(12). In this study, the reference strain of *T. asteroides* was isolated from the blood of patients with disseminated infections caused by *Trichosporon* spp.(11), which indicated high pathogenicity. *T. jirovecii* genotype 1 has been isolated from the human body(9), but its pathogenicity was unknown. Thus far, there have been few reports on *T. jirovecii:* Malgorzata et al. reported one case of mixed respiratory infection in a dog caused by *T. jirovecii* and *Rhodotorula*(31, 32), and Nardoni reported one case of back infection in a tortoise caused by *T. jirovecii*(7). In this study, four strains of *T. laibachii* (JYZ3252, JYZ921, JYZ321, and JYZ912) and one strain of *T. moniliiforme* (JYZ372) were identified as having new genotypes. Their pathogenicity remains to be confirmed by future studies.

### 4.3 Genotyping of *Trichosporon* spp.

At present, IGS1 sequence analysis is generally used for genotyping *Trichosporon* spp. For example, Chagasneto et al. completed the genotyping of 14 strains of *T. asahii* by IGS sequence analysis(11), whereas Guo completed the genotyping of 39 strains of *T. asahii*(8). The main target in the genotyping of *Trichosporon* spp. has been *T. asahii*, and the genotyping of other *Trichosporon* spp. has been rare. In this study, among all the isolates only *T. jirovecii* had been assigned to two genotypes, and the other species had not been studied(9). The main reason was that the identification of *Trichosporon* spp. is difficult and sequence information for IGS1 is scarce. In this study, only preliminary genotyping was performed for *Trichosporon* spp., and further research will rely on improvements in sequence information for IGS1in *Trichosporon* spp.

### 4.4 Morphological development process of *Trichosporon* spp.

There have been few studies on the morphology of *Trichosporon* spp. In 2005, Li et al. performed ITS-PCR detection and morphological and susceptibility testing on six *Trichosporon* spp.(13). The colonies of different *Trichosporon* spp. were similar, but the morphologies of their mycelia and spores were significantly different. The structure of the mycelium was not destroyed, and the test results were credible. The morphology of the mycelia of *T. domesticum* was very similar to that in this study. The morphological development process of *Trichosporon* spp. was significantly different, and the majority of *Trichosporon* spp. had a typical structure: for example, septal differentiation of the mycelium in *T. gracile* (Fig. 8D); short thick mycelium during the development of *T. shinodae* (Fig. 11C); elongation and bifurcation of the mycelium and the aggregation of spores into spheres in *T. asteroides* (as shown in Fig. 10C); and a spindle-type articular spore structure in *T. middelhovenii* (Fig. 13A, B, C, and D). The above results proved that the morphological development process and typical structure have great significance as references for morphological identification.

From the point of view of the development and sporulation of mycelia, *Trichosporon* is an intermediate genus between molds and yeasts. Its mycelia can differentiate into a large number of spores like yeasts and also produce conidia like molds. Spores in the early stages of development can either bud like hyphae or divide like those of yeasts. Colonies of some *Trichosporon* spp. resemble yeasts in being milky, oily, and reflective, whereas colonies of some *Trichosporon* spp. have a radiate texture similar to that of molds(13). *Trichosporon* might represent an intermediate genus in the evolution of yeasts into molds. In the study of the morphology of *Trichosporon* spp., they should be regarded as molds in order to observe their sporulation and mycelial structure.

### 4.5 Pathological changes in *Trichosporon* spp. infections

Different *Trichosporon* spp. cause similar pathological lesions on the skin and liver. *T. asahii* caused hepatic sinusoidal dilatation, mild to moderate dilatation of small blood vessels, hyperemia, neutrophil-based focal infiltration of inflammatory cells, and proliferation or degeneration of hepatocytes(33). *T. dermatis* caused hepatic sinusoidal dilatation and congestion, swelling, degeneration, or necrosis of hepatocytes, and hyperplasia of Kupffer cells(34). These lesions were similar to the pathological changes in the liver observed in this study. In the literature there are few mentions of skin lesions, subcutaneous abscesses, and bruises that were caused by *T. dermatis*.

However, most of the *Trichosporon* spp. identified in this study could cause skin lymphocyte infiltration, abscesses, and thickening of the stratum corneum.The pathological changes were significantly different between the groups treated by subcutaneous injection and skin inunction. These conclusions were similar to those of a study that was reported in China for the first time in 2010(35). The skin damage caused in the group treated by skin inunction was lighter, and only *T. laibachii* and *T. asteroides* caused obvious pathological changes, which might be related to the uncontrollable amount of the spore coating and the pathogenicity of the *Trichosporon* spp. themselves. The spores developed a strong tendency to form mycelium, and the process of formation of mycelium could cause mechanical damage, which might be a reason for these observations.

### 4.6 Pathogenicity of the *Trichosporon* spp.

Except for *T. moniliiforme, T. domesticum*, and *T. gracile*, all the *Trichosporon* spp. in this study caused significant damage to the liver and skin in healthy mice (Fig.S1~S10). In most cases spores stained with periodic acid/Schiff stain could be observed in skin sections. *T. brassicae, T. guehoae, T. middelhovenii*, and *T. shinodae* were found for the first time to be pathogenic forms of *Trichosporon* that could provoke obvious lesions in immunosuppressed and non-immunosuppressed groups. Hitherto, reports of the pathogenicity of these four *Trichosporon* spp. had not been found. This might be related to difficulties in the identification of *Trichosporon* spp. and differences in pathogenicity caused by the differences between strains.

In this study, *T. asteroides* (JYZ1255) exhibited strong pathogenicity. The infected tissue was extensively congested, and there was a large area of abscess. Two mice in group A died 2 days after being inoculated with a suspension of *T. asteroides*. There have been many reports on *T. asteroides*, which is one of the main pathogens involved in trichosporosis in humans and is a dominant strain among fungi on the body surface of the giant panda. *T. asteroides* caused purulent keratitis(36) and was also isolated from the blood of patients with disseminated trichosporosis(9, 11). *T. asteroides* gave rise to obvious disseminated infections in the reticular layer of the skin (Fig. 14G1) and budding in the dermis (Fig. 14G2). The infections might cause damage or even be life-threatening to immunocompromised giant pandas. *T. jirovecii* (JYZA10) was identified as having genotype 1 in previous studies (Fig. 1) and was significantly more pathogenic in immunosuppressed mice than in non-immunosuppressed mice and tissue. The degree of damage was significantly lower than that caused by *T. asteroides*, and it was inferred that genotype 1 of T *jirovecii* was opportunistically pathogenic. Four strains of *T. laibachii* were identified as having new genotypes by phylogenetic analysis of the IGS1 sequence (Fig. 2). Although there have currently been no reported cases of infection involving *T. laibachii, T. laibachii* (JYZ3252) caused skin ulceration in mice (Fig. 4). This lesion demonstrated that the new genotype of *T. laibachii* is more pathogenic.

The pathogenicity test only studied the effects of *Trichosporon* spp. on the skin and liver, and most of the isolated *Trichosporon* spp. caused more severe damage to skin than to the liver. For example, *T. brassicae* (JYZ1253) and the reference strain, which was isolated from rancid milk, had the same genotype, but the isolated strain caused severe inflammatory reactions in skin tissue and skin necrosis. *T. asteroides* (JYZ1255) and the reference strain, which was isolated from human blood, had the same genotype, but the isolated strain caused disseminated infections in the reticular layer of the skin. It was assumed that the pathogenicity of *Trichosporon* spp. is related to their parasitic environment and that *Trichosporon* spp. that are isolated from the skin surface cause more pronounced damage to the skin.

## List of abbreviations

There are no abbreviations.

## Declarations

### Ethics approval and consent to participants

All animal experiments were approved by the Institutional Animal Care and Use Committee of the Sichuan Agricultural University (permit number DYY-S20151326). Permissions were obtained from the China Conservation and Research Center for the Giant Panda Breeding prior to sample collection from the pandas.

### Consent for publication

Not applicable

### Authors' contributions

XM, YJ,CW,YG,SC,XH YW,QZ,RW,QY and XH carried out the collection of the sample of the giant pandas, isolated fungi, conceived the study, and drafted the manuscript. ZZ, JD, ZR, SY, LS and GP participated in the sequence alignment, carried out the molecular genetic studies, and participated in the data analysis. HL and ZZconceived the study, participated in its design and coordination, and helped draft the manuscript. All authors read and approved the final manuscript.

